# Timing of selective basal ganglia white matter loss in Huntington’s disease

**DOI:** 10.1101/2021.02.17.431568

**Authors:** Paul Zeun, Peter McColgan, Thijs Dhollander, Sarah Gregory, Eileanoir B Johnson, Marina Papoutsi, Akshay Nair, Rachael I Scahill, Geraint Rees, Sarah J Tabrizi, the TrackOn-HD and HD-YAS Investigators

**Affiliations:** Huntington’s Disease Centre, Department of Neurodegenerative disease, UCL Queen Square Institute of Neurology, University College London, WC1N 3BG, UK; The Murdoch Children’s Research Institute, Parkville Victoria 3052, Australia; Max Planck UCL Centre for Computational Psychiatry and Ageing Research, UCL Queen Square Institute of Neurology, University College London, WC1N 3BG, UK; UCL Institute of Cognitive Neuroscience, Queen Square, London WC1N 3BG, UK; Dementia Research Institute at UCL, London, WC1N 3BG, UK

## Abstract

**Objectives:** To investigate the timeframe prior to symptom onset when cortico-basal ganglia white matter (WM) loss begins in premanifest Huntington’s disease (preHD), and which striatal and thalamic sub-region WM tracts are most vulnerable.

**Methods:** We performed fixel-based analysis, which allows resolution of crossing WM fibres at the voxel level, on diffusion tractography derived WM tracts of striatal and thalamic sub-regions in two independent cohorts; TrackON-HD, which included 72 preHD (approx. 11 years before disease onset) and 85 controls imaged at three time points over two years; and the HD young adult study (HD-YAS), which included 54 preHD (approx. 25 years before disease onset) and 53 controls, imaged at one time point. Group differences in fibre density and cross section (FDC) were investigated.

**Results:** We found no significant group differences in cortico-basal ganglia sub-region FDC in preHD gene carriers 25 years before onset. In gene carriers 11 years before onset, there were reductions in striatal (limbic and caudal motor) and thalamic (premotor, motor and sensory) FDC at baseline, with no significant change over 2 years. Caudal motor-striatal, pre-motor-thalamic, and primary motor-thalamic FDC at baseline, showed significant correlations with the Unified Huntington’s disease rating scale (UHDRS) total motor score (TMS). Limbic cortico-striatal FDC and apathy were also significantly correlated.

**Conclusions:** Our findings suggest that the initiation of disease modifying therapies 25 years before onset could protect these important brain networks from undergoing neurodegeneration and highlight selectively vulnerable sub-regions of the striatum and thalamus that may be important targets for future therapies.

## Introduction

Huntington’s disease (HD) is a neurodegenerative condition caused by a trinucleotide repeat expansion in the huntingtin gene. This results in degeneration of the cortico-basal ganglia white matter (WM) circuits resulting in progressive motor, cognitive and neuropsychiatric disturbance^1^ ^2^.

The basal-ganglia shows some of the earliest changes in premanifest (preHD), with loss of striatal grey matter (GM)^3^ and WM^4^ ^5^ and posterior WM thalamic radiations^6^ occurring 15 years before disease onset.

In manifest HD diffusion tractography^7^ ^8^ and anatomically based parcellations^9^ of striatal GM show group differences in cognitive and motor sub-regions. In preHD anterior caudate - frontal eye field connectivity has been associated with deficits in saccadic eye movements^10^. A study from PREDICT-HD has investigated the WM tracts of motor and sensory striatal sub-regions and demonstrated widespread group differences at baseline and change in the premotor striatum WM over time^11^.

To date no study has investigated WM tracts of all functional striatal and thalamic sub-regions in HD. Gene therapy trials in HD^12^ are targeting selective basal ganglia sub-regions, therefore understanding selective sub-region vulnerability is vital to designing these therapies.

Here we aim to address two questions: which striatal and thalamic sub-regions WM tracts are most vulnerable in preHD and what is the time frame prior to symptom onset when these changes begin. In order to do this we performed diffusion MRI (dMRI) fixel-based analysis in two independent preHD cohorts; TrackON-HD^13^ (approx. 15 yrs before disease onset) and the HD young adult study (HD-YAS)^14^ (approx. 25 years before disease onset).

## Methods

An overview of study methodology and key results is provided in figure 1. For each cohort, gene carriers were required to have no diagnostic motor features of the disease (Unified Huntington’s disease rating scale (UHDRS) total motor score (TMS) diagnostic confidence level (DCL) of <4) and a CAG repeat size of ≥40 where there is complete disease penetrance. Controls were either gene negative (family history of HD but negative genetic test), partners of gene carriers, or members of the wider HD community (recruited through support groups or friends of participants). Participants were excluded if they were left-handed, ambidextrous, or had poor quality diffusion MRI data, as defined by visual quality control performed by PM and SG (Supplementary table S1). Additional exclusion criteria included age below 18 or above 65, major psychiatric, neurological or medical disorder or a history of severe head injury. See Kloppel et al.^13^ and Scahill et al.^14^ for detailed criteria. Participant demographics are summarised in supplementary table S2. HD-YAS preHD gene carriers were an average of 25 years before predicted disease onset, while Track-ON preHD gene carriers were an average of 11 years before disease onset. Year before predicted disease onset was calculated using the Langbehn equation^15^. This is based on a parametric survival model developed using 2,913 HD gene carriers from 40 centres worldwide. Both studies were approved by the local ethics committee and all participants gave written informed consent according to the Declaration of Helsinki.

**Figure 1.**
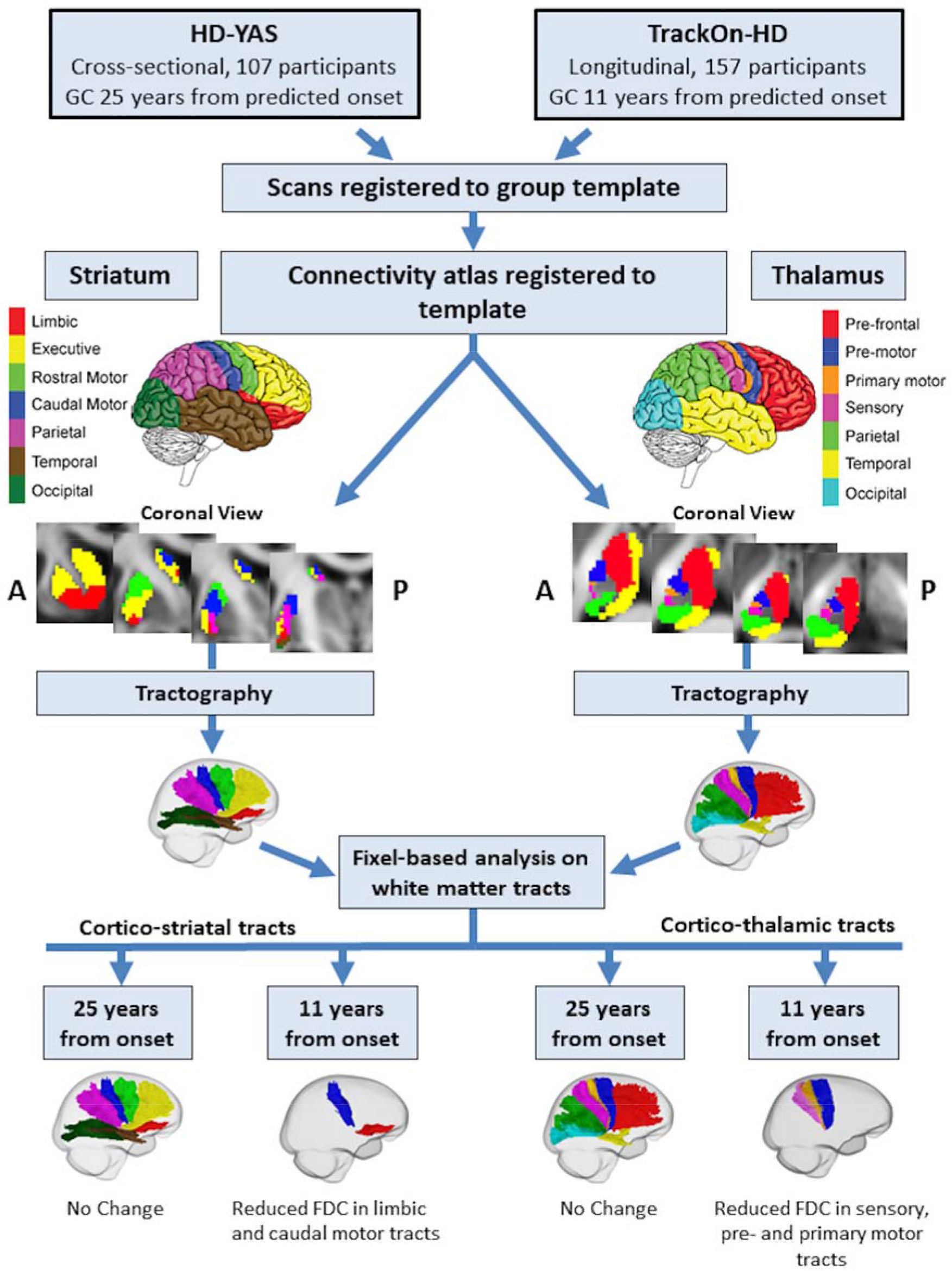
Overview of study methodology and key results. Diffusion data were analysed from two separate studies; HD-YAS, where HD gene carriers were 25 years before predicted disease onset, and TrackON-HD, where HD gene carriers were 11 years before predicted onset. For each study, scans were registered to a common template. Connectivity based atlases of the striatum and thalamus were registered to the group template. Diffusion tractography was performed on the group template to reconstruct each of the cortico-thalamic and cortico-striatal tracts in right and left hemispheres. Measures of FDC were then computed for each tract. In gene carriers 25 years before predicted onset, there were no differences in any cortico-striatal or cortico-thalamic tract. In HD gene carriers 11 years from predicted onset, we found reduced FDC in limbic and caudal motor cortico-striatal tracts and pre-motor, primary motor and sensory cortico-thalamic tracts cross-sectionally. There were no significant longitudinal changes in gene carriers 11 years before predicted onset. GC = Gene Carriers, FDC = Fibre density and cross-section.

### HD-Young Adult Study

To investigate how early basal ganglia white matter loss could be detected, we utilised MRI data from the HD-young adult study (HD-YAS).^16^ This was a single site cross-sectional study of gene carriers and controls aged 18-40. Data were collected between the August 2017 and April 2019. PreHD gene carriers required a disease burden score, a product of age and CAG repeat length^17^ of ≤ 240 indicating these participants were ≥ 18 years before predicted disease onset. Multi-shell dMRI data were analysed from 54 preHD and 53 control participants.

### TrackON-HD study

To investigate whether there is selective loss of specific basal ganglia white matter connections, we utilised MRI data from the TrackOn-HD study^18^. This included single-shell diffusion data from preHD and control participants, scanned at 3 time-points one year apart over 2 years from 4 sites (London, Paris, Leiden, and Vancouver). Gene carriers were required to have a disease burden score of > 250 indicating participants were < 17 years before predicted onset. The total number of participants each year was as follows: year 1 (72 gene carriers, 85 controls), year 2 (81 gene carriers, 87 controls) and year 3 (80 gene carriers, 78 controls).

At the last TrackON-HD visit, participants at London and Paris sites had an additional multi-shell dMRI scan which included 33 gene carriers and 40 healthy control participants (Supplementary table S2).

### MRI acquisition

For the HD-YAS acquisition, dMRI data were acquired using a 3T Siemens Prisma scanner, with a 64-channel head coil and b-values of 0, 300, 1000 and 2000 s/mm^2^ with 10, 8, 64 and 64-gradient directions respectively. Images had a voxel size of 2×2×2 mm^3^, 72 slices and a repetition time/time of echo of 3260/58 ms. One of the b=0 volumes was acquired with reverse phase-encoding to allow for susceptibility-induced distortion correction. The scanning time was 10 minutes.

TrackON-HD single-shell dMRI were acquired using 4 different 3T scanners (Siemens TIM Trio MRI scanners at London and Paris, Phillips Achieva at Vancouver and Leiden) using similar imaging acquisitions across sites. Using a 12-channel head coil, dMRI were acquired with 42 unique gradient directions (*b* = 1000 s/mm^2^). Eight and one images with no diffusion weighting (*b* = 0) were acquired using the Siemens and Philips scanners, respectively. For the Siemens and Phillips scanners, repetition time/time of echo was 1300/88 and 1100/56ms respectively. Voxel size for the Siemens scanners was 2×2×2 mm^3^, and voxel size for the Phillips scanners was 1.96×1.96×2 mm^3^. 75 slices were collected for each diffusion-weighted and non-diffusion-weighted volume with a scanning time of 10 minutes.

In additional to single shell dMRI, for the final time point in TrackON-HD, multi-shell dMRI data were also acquired at London and Paris, with b =300, 700, 2000 sec/mm^2^ and 8, 32 and 64 gradient directions respectively; 14 b=0 images; voxel size = 2.5×2.5×2.5 mm^3^; repetition time/time of echo = 7000 ms/ 90.8 ms; 55 slices and a scanning time 15 minutes (Supplementary table S2). Multi-shell and single shell acquisitions were analysed separated and study site was included as a covariate in all analyses.

### Diffusion MRI processing

Preprocessing of dMRI was performed using MRtrix3^19^ and FSL^20^ software packages. This included denoising of data^21^, Gibbs-ringing artefact removal^22^, eddy-current correction and motion correction^23^, and up-sampling diffusion MRI spatial resolution in all 3 dimensions using cubic b-spline interpolation to 1.3×1.3×1.3 mm^3^ voxels^24^. The upsampling of data helps to increase the anatomical contrast, which improves downstream spatial normalisation and statistics^25^.

Three-tissue constrained spherical deconvolution (CSD) modelling of diffusion data was performed using MRtrix3Tissue (https://3Tissue.github.io), a fork of MRtrix3. For all data, response functions for single-fibre white matter as well as GM and CSF were estimated from the data themselves using an unsupervised method^26^. For the single-shell TrackON-HD data, WM-like fibre orientation distributions (FODs) as well as GM-like and CSF-like compartments in all voxels were then computed using Single-Shell 3-Tissue CSD^27^, whilst multi-shell multi-tissue CSD^28^ was utilised for the TrackON-HD multi-shell and HD-YAS data. Spatial correspondence was achieved by generating a group-specific population template with an iterative registration and averaging approach using white matter FOD images for 40 subjects (20 preHD and 20 controls, selected at random), in keeping with previous studies^29^. Each subject’s FOD image was then registered to the template via a FOD-guided non-linear registration^30^. For longitudinal data, an intra-subject template was produced using scans from all 3-time points before creating a common population template using the same registration approach.

### Fixel based analysis

We utilised a fixel-based analysis to interrogate changes in white matter for this analysis. This was implemented in MRtrix3^19^ and the methodology has been described previously^25^. In brief, fibre density (FD) is a measure of the intra-axonal volume of white matter axons aligned in a particular direction^31^. The fibre bundle cross section (FC) measures the cross-section of a fibre bundle by using the non-linear warps required to spatially normalise the subject image to the template image. The fibre density and cross section (FDC) is calculated by multiplying FD and FC providing one metric sensitive to both fibre density and fibre cross-section. For the main analysis, we report FDC only as it can be expected that neurodegeneration will cause combined reductions in both fibre density and bundle atrophy, however specific FD and FC measures are reported in the supplementary tables.

### Generating tracts for analysis

We reconstructed distinct cortico-striatal and cortico-thalamic tracts using diffusion tractography on a group template before comparing FDC for each tract between groups (Fig. 1). Atlases of the striatum and thalamus derived using diffusion tractography were used to segment each structure in to 7 sub-regions per hemisphere based on the dominant area of cortical connectivity for each sub-region. For the striatum, the 7 sub-regions included limbic, executive, rostral motor, caudal motor, parietal, temporal and occipital regions^32^. The limbic sub-region connects to the orbital gyri, gyrus rectus and ventral anterior cingulate. The executive tract connects to dorsal prefrontal cortex. The rostral motor area connects to the pre-supplementary motor cortex and the frontal eye field region. The caudal motor tract connects the post-commissural striatum to the primary motor cortex, whilst parietal, temporal and occipital sub-regions connect to respective cortices. For the thalamic segmentation, the seven sub-regions are defined as those connected to prefrontal, premotor, primary motor, sensory motor, parietal, temporal and occipital cortices^33^. Within this segmentation, medial and dorsal thalamus including the mediodorsal nucleus connects to prefrontal and temporal regions. The ventral posterior nucleus connects to sensory motor cortex. The ventral lateral and anterior nuclei connect to primary motor and premotor cortex. The lateral posterior nucleus and parts of the pulvinar connect to parietal cortex.

The striatal and thalamic atlases were registered to the population template using NiftyReg^34^ using a linear registration. Registration for each subject was visually checked to ensure accurate registration. To generate a tract for each striatal and thalamic sub-region, a tractogram was generated using probabilistic tractography on the population template. 20,000 streamlines were generated for each individual tract. Streamlines were initiated in each sub-region, with all other sub-regions masked out to avoid streamlines traversing other sub-regions and creating large amounts of overlap between the tracts. The result was a single fibre bundle for each sub-region connecting to its respective main cortical region. Fixel-based metrics were then calculated for all fixels (analogous to voxels) across the entire tract and averaged to generate a single measure of FDC for each tract. While mild striatal and cortical grey matter atrophy occurs in preHD, we did not perform volume normalisation as this can over compensate^35^ leading to spurious results^4^.

### Statistical analysis

All statistical analysis was performed in MATLAB R2018a. Statistical analysis of the single-time point HD-YAS and TrackON-HD multi-shell data involved permutation testing (10,000 permutations) with two-tailed t-tests to investigate group differences. Age, gender, study site and education were included as covariates.

For the longitudinal Track-On analysis, linear mixed effects regression (LMER) was used, as it provides unbiased estimates under the assumption that the missing data is ignorable. It also accounts for dependence due to repeated measures. This approach is similar to previous longitudinal HD imaging studies^5^ ^36^. The LMER model is defined as:

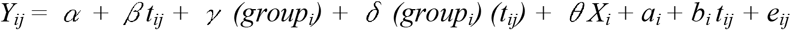

Where Y_*ij*_ is FDC for the *i^th^* participant ( *i = 1, …, N*) at the *j^th^* time point (*j = 1, …, ni*), with time metric *t_ij_ = visit_ij_ – 1*, so that *t_i1_ = 0* . Furthermore, *group_i_* is a dummy variable taking the value of 0 if a participant is in the control group and the value of 1 if a gene carrier.

Greek letters denote fixed effects; *α* is the control group mean at the first visit, *β* is the control group linear slope, *γ* is the mean difference among the preHD and control groups at the first visit (difference of intercepts), *δ* is the slope difference among the groups (rate of change difference), *X_i_* is the matrix of covariates (age, sex, site, education) with associated regression coefficient vector *θ*; *a_i_* and *b_i_* are random effects (random intercepts and slopes), and *e_ij_* is random error. Maximum likelihood methods are used for estimation under the assumption that the random effects have a joint-normal distribution with zero-means and non-singular covariance matrix, and the random error is normally distributed with zero mean and constant nonzero variance. The objects of inference were *γ* and *δ*, with the former being the initial cross-sectional mean difference among the groups adjusting for the covariates and the latter being the group difference in the rate of change (slope difference) adjusting for the covariates. The null hypothesis of interest were *H_0_: γ* = *0* (no initial mean group difference) and *H_0_*: *δ* = *0* (no group difference in rate of change), which were tested with the *z*-values of *z* = *γ □*/*SE(γ* □) and *z* = *δ* □ / *SE (δ* □).

We addressed multiple comparisons by applying a false discovery rate (FDR) approach to each separate cohort analysis and considered an FDR estimate of ≤ 0.05 to be significant.

To investigate whether changes in FDC show a relationship with clinical measures, a priori clinical correlations for preHD in the TrackON-HD (n = 72), were performed for the tracts showing baseline change in the TrackON-HD single-shell results. Apathy scores from the Baltimore Apathy and Irritability scale^37^ (10.8 ± 7.4) were selected for limbic cortico-striatal tracts and the UHDRS^38^ TMS (5.9 ± 3.7) and DCL (1.4 ± 1) for caudal motor cortico-striatal tracts and thalamic tracts to premotor and primary motor cortices. Correlations were performed using partial correlations with age, gender, site, education and CAG included as covariates.

### Data Availability

The data that support the findings of this study are available from the corresponding author upon reasonable request.

## Results

### No significant differences in cortico-striatal and cortico-thalamic connections 25 years before predicted onset in HD gene carriers

In the group of HD gene carriers approximately 25 years before predicted onset, no significant changes were seen in any cortico-striatal or cortico-thalamic tract compared to matched controls after FDR correction (Fig. 2 A-B and 3 A-B and tables 1–2). This suggests that cortico-basal ganglia white matter connections 25 years before predicted disease onset are structurally preserved.

**Figure 2.**
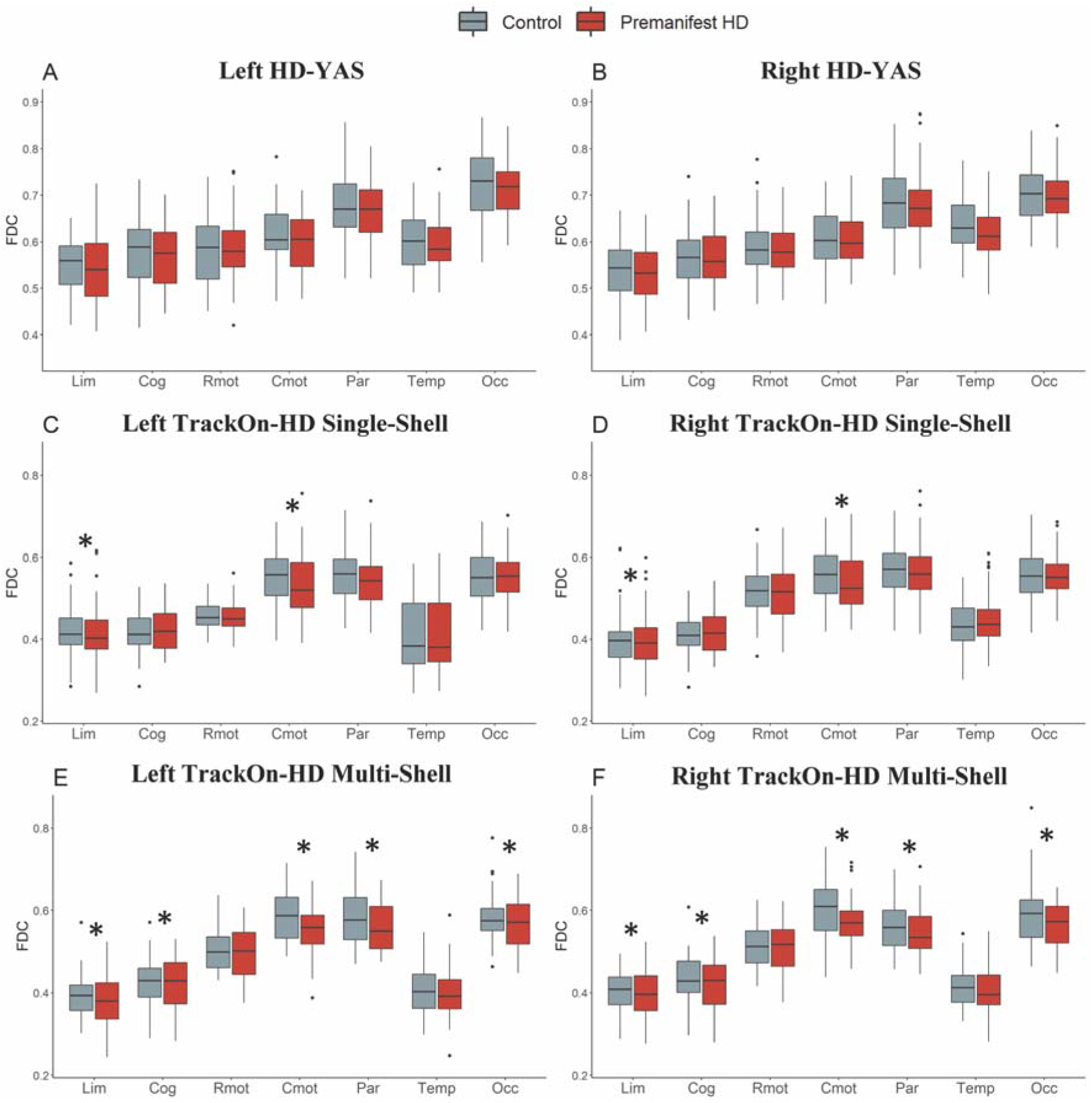
Cortico-striatal tract fibre density and cross-section left and right in HD-YAS (A+B), TrackOn-HD single-shell (B+C) and multi-shell (E+F) datasets. FDC = Fibre density and cross-section. Cortico-striatal tracts are displayed on the x-axis. Lim = Limbic, Cog = Cognitive, Rmot = Rostral motor, Cmot = Caudal motor, Par = Parietal, Temp = Temporal, Occ = Occipital. * FDR= < 0.05.

**Figure 3.**
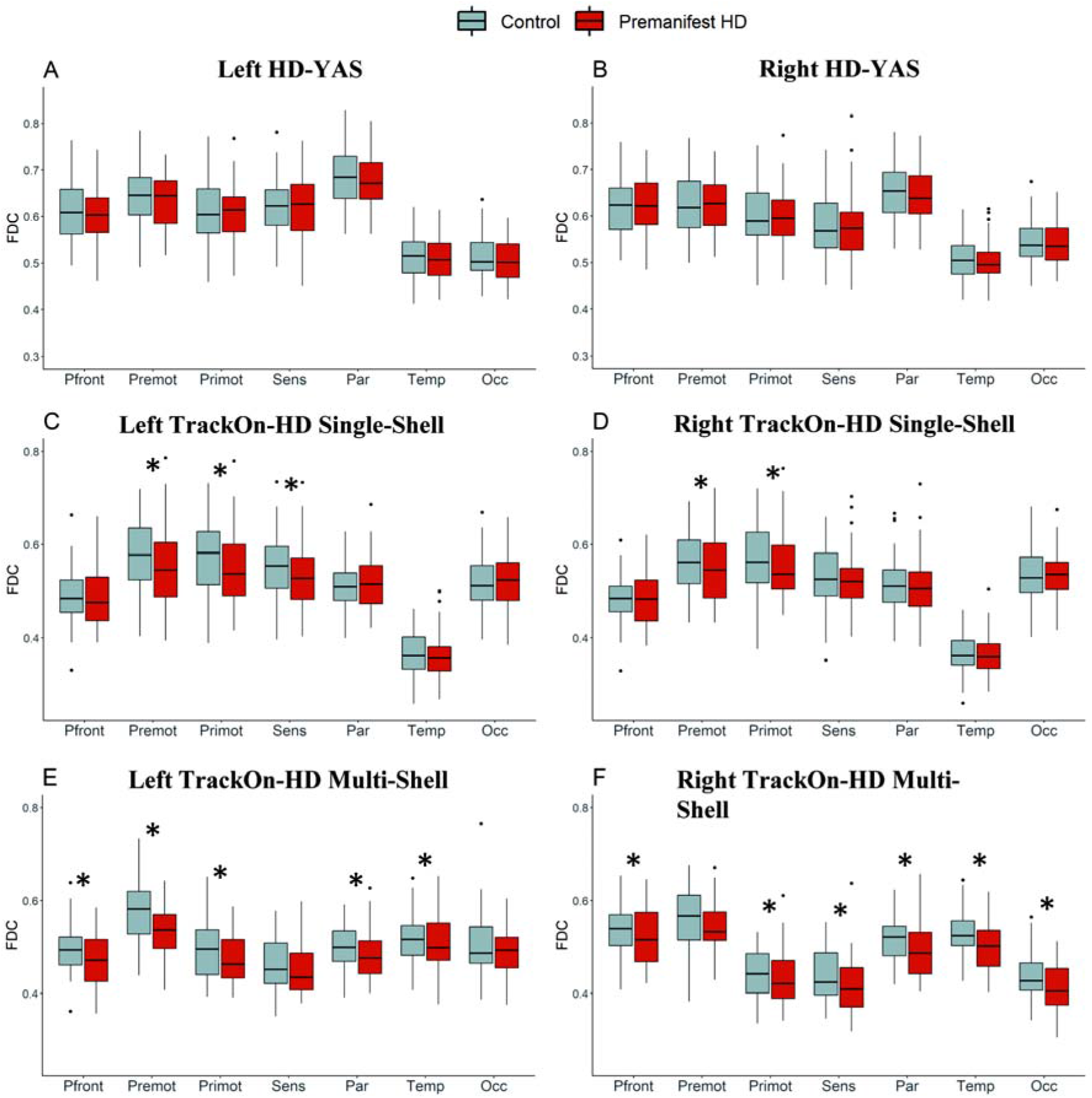
Cortico-thalamic tract fibre density and cross-section left and right in HD-YAS (A+B), TrackOn-HD single-shell (C+D) and multi-shell (E+F) datasets. FDC = Fibre density and cross-section. Cortico-thalamic tracts are displayed on the x-axis. Pfront = Pre-frontal, Premot = Pre-motor, Primot = Primary motor, Sens = Sensory, Par = Parietal, Temp = Temporal, Occ = Occipital. * FDR = < 0.05.

**Table 1.** Cortico-striatal fibre density and cross section (FDC) in HD-YAS

| Cortico-striatal Tract | Control Mean | PreHD Mean | SE | p | FDR |
| --- | --- | --- | --- | --- | --- |
| **L Limbic** | 0.55 | 0.55 | 0.001 | 0.31 | 0.38 |
| **R Limbic** | 0.54 | 0.53 | 0.001 | 0.16 | 0.35 |
| **L Cognitive** | 0.58 | 0.57 | 0.001 | 0.32 | 0.38 |
| **R Cognitive** | 0.56 | 0.57 | 0.001 | 0.49 | 0.53 |
| **L Rostral Motor** | 0.58 | 0.59 | 0.001 | 0.54 | 0.54 |
| **R Rostral Motor** | 0.59 | 0.59 | 0.001 | 0.33 | 0.38 |
| **L Caudal Motor** | 0.62 | 0.60 | 0.001 | 0.04 | 0.35 |
| **R Caudal Motor** | 0.61 | 0.60 | 0.001 | 0.16 | 0.35 |
| **L Parietal** | 0.67 | 0.67 | 0.001 | 0.33 | 0.38 |
| **R Parietal** | 0.68 | 0.68 | 0.001 | 0.34 | 0.38 |
| **L Temporal** | 0.60 | 0.60 | 0.001 | 0.18 | 0.35 |
| **R Temporal** | 0.64 | 0.61 | 0.001 | 0.01 | 0.22 |
| **L Occipital** | 0.73 | 0.71 | 0.001 | 0.07 | 0.35 |
| **R Occipital** | 0.71 | 0.70 | 0.001 | 0.18 | 0.35 |

Unadjusted means displayed. $\gamma$ =estimated group intercept difference, SE=standard error, p = p-value, FDR = false discovery rate corrected p-value.

**Table 2.** Cortico-thalamic fibre density and cross section (FDC) in HD-YAS

| Cortico-thalamic Tract | Control Mean | PreHD Mean | SE | p | FDR |
| --- | --- | --- | --- | --- | --- |
| **L Prefrontal** | 0.61 | 0.61 | 0.001 | 0.18 | 0.35 |
| **R Prefrontal** | 0.62 | 0.62 | 0.001 | 0.32 | 0.38 |
| **L Premotor** | 0.65 | 0.63 | 0.001 | 0.08 | 0.35 |
| **R Premotor** | 0.63 | 0.62 | 0.001 | 0.30 | 0.35 |
| **L Primary Motor** | 0.61 | 0.61 | 0.001 | 0.34 | 0.38 |
| **R Primary Motor** | 0.60 | 0.60 | 0.001 | 0.34 | 0.38 |
| **L Sensory** | 0.62 | 0.62 | 0.001 | 0.52 | 0.54 |
| **R Sensory** | 0.58 | 0.58 | 0.001 | 0.33 | 0.38 |
| **L Parietal** | 0.68 | 0.67 | 0.001 | 0.09 | 0.35 |
| **R Parietal** | 0.65 | 0.64 | 0.001 | 0.17 | 0.35 |
| **L Temporal** | 0.52 | 0.51 | 0.0004 | 0.13 | 0.35 |
| **R Temporal** | 0.51 | 0.50 | 0.0004 | 0.05 | 0.35 |
| **L Occipital** | 0.51 | 0.50 | 0.0004 | 0.08 | 0.35 |
| **R Occipital** | 0.55 | 0.54 | 0.0004 | 0.18 | 0.35 |

Unadjusted means displayed. $\gamma$ =estimated group intercept difference, SE=standard error, p = p-value, FDR = false discovery rate corrected p-value.

### Anatomically specific basal ganglia white matter loss in preHD

We applied the same technique to a cohort of HD gene carriers closer to predicted onset to establish whether particular connections showed selective loss at this stage using the TrackON-HD single-shell cohort baseline results. For cortico-striatal tracts, significant reductions were seen bilaterally in limbic (left FDR = 0.002, right FDR = 0.02) and caudal motor (left FDR = 0.002, right FDR = 0.006) FDC (Fig. 2 C-D and table 3). For cortico-thalamic tracts, significant reductions were seen bilaterally in pre-motor (left FDR = 0.005, right FDR = 0.02), primary motor (left FDR = 0.001, right FDR = 0.02) and left sensory (FDR = 0.006) FDC (Fig. 3 C-D and table 4).

**Table 3.** Cortico-striatal fibre density and cross section (FDC) in TrackOn-HD Single-Shell Baseline

| **Cortico-striatal Tract** | **Control Mean** | **PreHD Mean** | **$\gamma$** | **SE** | **p** | **FDR** |
| --- | --- | --- | --- | --- | --- | --- |
| **L Limbic** | 0.42 | 0.41 | -0.03 | 0.008 | $3.0 \times 10^{-4}$ | 0.002 |
| **R Limbic** | 0.40 | 0.39 | -0.02 | 0.007 | 0.005 | 0.02 |
| **L Cognitive** | 0.42 | 0.42 | -0.01 | 0.006 | 0.05 | 0.08 |
| **R Cognitive** | 0.41 | 0.42 | -0.01 | 0.006 | 0.13 | 0.15 |
| **L Rostral Motor** | 0.49 | 0.50 | -0.007 | 0.008 | 0.38 | 0.55 |
| **R Rostral Motor** | 0.52 | 0.52 | -0.02 | 0.008 | 0.04 | 0.09 |
| **L Caudal Motor** | 0.55 | 0.53 | -0.03 | 0.008 | $4.9 \times 10^{-4}$ | 0.002 |
| **R Caudal Motor** | 0.56 | 0.54 | -0.03 | 0.008 | 0.001 | 0.006 |
| **L Parietal** | 0.55 | 0.55 | -0.02 | 0.008 | 0.02 | 0.06 |
| **R Parietal** | 0.57 | 0.56 | -0.02 | 0.008 | 0.07 | 0.11 |
| **L Temporal** | 0.41 | 0.41 | $2.2 \times 10^{-4}$ | 0.006 | 0.97 | 0.97 |
| **R Temporal** | 0.43 | 0.44 | 0.001 | 0.006 | 0.84 | 0.84 |
| **L Occipital** | 0.55 | 0.55 | -0.004 | 0.007 | 0.53 | 0.62 |
| **R Occipital** | 0.56 | 0.56 | -0.01 | 0.007 | 0.08 | 0.11 |

Unadjusted means displayed. $\gamma$ =estimated group intercept difference, SE=standard error, p = p-value, FDR = false discovery rate corrected p-value.

**Table 4.** Cortico-thalamic fibre density and cross section (FDC) in TrackOn-HD Single-Shell Baseline cohort

| **Cortico-thalamic Tract** | **Control Mean** | **PreHD Mean** | **$\gamma$** | **SE** | **p** | **FDR** |
| --- | --- | --- | --- | --- | --- | --- |
| **L Prefrontal** | 0.49 | 0.49 | -0.02 | 0.007 | 0.03 | 0.05 |
| **R Prefrontal** | 0.48 | 0.48 | -0.01 | 0.007 | 0.08 | 0.08 |
| **L Premotor** | 0.58 | 0.55 | -0.03 | 0.009 | 0.002 | 0.005 |
| **R Premotor** | 0.51 | 0.51 | -0.03 | 0.009 | 0.002 | 0.02 |
| **L Primary Motor** | 0.57 | 0.55 | -0.03 | 0.008 | $1.30 \times 10^{-4}$ | 0.001 |
| **R Primary Motor** | 0.57 | 0.55 | -0.02 | 0.009 | 0.007 | 0.02 |
| **L Sensory** | 0.55 | 0.53 | -0.02 | 0.008 | 0.003 | 0.006 |
| **R Sensory** | 0.53 | 0.52 | -0.01 | 0.007 | 0.08 | 0.08 |
| **L Parietal** | 0.51 | 0.51 | -0.006 | 0.006 | 0.35 | 0.40 |
| **R Parietal** | 0.51 | 0.51 | -0.01 | 0.006 | 0.05 | 0.08 |
| **L Temporal** | 0.36 | 0.36 | -0.007 | 0.005 | 0.15 | 0.21 |
| **R Temporal** | 0.36 | 0.36 | -0.008 | 0.004 | 0.06 | 0.08 |
| **L Occipital** | 0.52 | 0.52 | -0.002 | 0.007 | 0.08 | 0.35 |
| **R Occipital** | 0.54 | 0.53 | -0.01 | 0.008 | 0.06 | 0.08 |

Unadjusted means displayed. $\gamma$ =estimated group intercept difference, SE=standard error, p = p-value, FDR = false discovery rate corrected p-value.

We next investigated whether there were significant changes over a 2-year time period at this stage of the disease. No significant changes in any cortico-striatal or cortico-thalamic tracts were seen after FDR correction (Supplementary tables S3-4).

### FDC changes using multi-shell acquisition at last time point in TrackOn-HD

We investigated the impact of dMRI acquisition on these results by repeating the analysis using the existence of an additional multi-shell dMRI scan, similar to the HD-YAS acquisition, in a subgroup of participants at the last time point in TrackON-HD. In this subgroup, we found widespread FDC changes in cortico-striatal and cortico-thalamic tracts (Fig. 2 E-F and 3 E-F and supplementary tables S5-6). Changes in FD and FC are presented in supplementary tables S7-12.

### Reductions in FDC correlate with a priori clinical measures

We assessed whether TrackON-HD baseline FDC changes were associated with relevant clinical measures in preHD (Supplementary table S13). FDC in caudal motor-striatal tracts significantly correlated with UHDRS TMS (left r = −0.22, p = 0.007, right r = −0.21, p = 0.009) after adjustment for age, sex, site and education. There were also significant correlations between UHDRS TMS and pre-motor-thalamic FDC (left r = −0.21, p = 0.01, right r = −0.20, p = 0.01) and primary motor-thalamic FDC (left r = −0.23, p = 0.004, right r = −0.20, p = 0.01). There were significant correlations between limbic cortico-striatal FDC and apathy (left r = 0.15, p = 0.07, right r = 0.22, p = 0.006).

## Discussion

By studying two unique cohorts, we provide new insights into the time frame and anatomical specificity of basal ganglia white matter loss in preHD. We show that cortico-striatal and cortico-thalamic WM tracts appear structurally preserved in preHD approximately 25 years before clinical onset. In a group approximately 11 years before clinical onset, specific cortico-striatal and cortico-thalamic tracts are more vulnerable than others to early degeneration, which is associated with emerging clinical signs. Thus the onset of basal ganglia WM loss is likely to occur in the time frame of 11-25 years before predicted onset. This is an important finding as it advances our understanding of the temporal pathology of HD beyond post-mortem studies. A follow-up study is planned in order to narrow this timeframe further and precisely define the optimal time-point for therapeutic intervention.

The striatum and thalamus have a distinct topographical organisation of cortical white matter tracts forming individual subnetworks^32^ ^33^. To our knowledge, this is the first time that preserved WM tracts across all striatal and thalamic sub-regions have been demonstrated in preHD. Recent evidence has suggested that the HD mutation may lead to abnormalities in striatal and cortical development^39^ ^40^. Our findings indicate that there is no detectable developmental abnormality in the microstructure of cortico-striatal and cortico-thalamic white matter tracts in preHD, using novel fixel based dMRI analysis, which has been shown to be more sensitive to neurodegeneration of WM in Parkinson’s^41^ and Alzheimer’s disease ^29^ when compared to standard DTI approaches.

Our findings suggest the initiation of disease modifying therapies very early in the premanifest period could help preserve these essential WM tracts, which form the motor, cognitive and limbic cortico-basal ganglia circuitry. Furthermore, delivery of emerging viral-vector based therapeutics such as RNAi^42^ by injection into the striatum and thalamus within this timeframe may allow for widespread drug distribution in the brain via persevered WM tracts to the cortex.

By using detailed tractography-based striatum and thalamus connectivity atlases, we show that specific basal ganglia sub-region WM tracts are more susceptible in preHD. Previous studies have investigated cortico-striatal WM tracts to all cortical regions only in manifest disease, where widespread group differences are already apparent across numerous WM tracts.^7^ ^8^ The only previous study in preHD assessing specific WM tracts of the striatum focused on motor, premotor and primary sensory WM tracts only, reporting cross-sectional differences largely in the group < 8 years before predicted onset^11^. Such previous studies also focus on DTI metrics, which are difficult to interpret in brain areas with crossing white matter fibres and have poor biological interpretability. In this study we apply fixel-based analysis, which is capable at resolving crossing WM fibres at the voxel level^25^. We present the first analyses to investigate all WM tracts of striatal sub-regions in preHD and in doing so identify the caudal motor-striatal and limbic-striatal WM tracts as being selectively vulnerable.

Whilst the striatum has long been an important target for disease-modifying therapies, the thalamus is one of the major connection hubs in the human cortex and as such may represent a target for RNAi injections to achieve widespread cortico-striatal coverage^43^. This study is the first to investigate thalamic sub-region WM tracts in HD. We show that thalamic WM tracts are persevered 25 years before disease onset in the HD-YAS cohort, however 11 years before disease onset there is selective vulnerability of pre-motor, primary motor and sensory thalamic WM tracts. This is in keeping with post-mortem evidence of thalamic selective vulnerability, where selective degeneration of the motor ventrolateral nucleus and the centromedian nucleus, which helps integrate sensorimotor functions, has been observed^44^.

The clinical relevance of this basal-ganglia WM tract selective vulnerability is demonstrated by negative correlations between the UHDRS TMS and FDC of the caudal motor-striatal, premotor-thalamic and primary motor-thalamic WM tracts, such that lower FDC is associated with greater motor deficit. The association of thalamic WM tracts with motor deficits suggests that in addition to the striatum, the thalamus may also prove to be an important therapeutic target to prevent early neurodegeneration and emerging motor symptoms. The lack of significant change in our longitudinal analysis indicates WM changes occur slowly over time in preHD, consistent with previous findings^45^.

A positive correlation is seen between limbic-striatal FDC and apathy, suggesting greater fibre density and fibre cross-section of the limbic-striatal WM tract is associated with greater apathy. While this may seem counter intuitive we have previously demonstrated positive association between apathy and functional connectivity of limbic regions including the anterior cingulate and medial orbitofrontal gyri^46^. This suggests apathy may be related to a hyper-connectivity state, with greater limbic-striatal FDC demonstrated here in keeping with the corresponding structural substrate of increased functional connectivity.

This is the first application of a fixel-based analysis in HD. The majority of previous dMRI studies in HD have been performed using the DTI model for tractogram generation and interpretation of WM microstructure. The DTI model has major limitations in its interpretability since most WM voxels contain contributions from multiple fibre populations and therefore voxel-averaged DTI measures are not fibre specific and have poor interpretability as measures of WM connectivity^25^ ^47^. Furthermore, many previous dMRI studies in HD^48^ have used a tract-based spatial statistics approach, which skeletonises the WM, but in doing so is unable to study specific WM tracts of interest, as done here. Fixel-based analysis accounts for crossing fibre populations to provide more reliable tractograms and account for the differing ways in which changes to intra-axonal volume may occur by quantifying both a measure of fibre density and a measure of fibre bundle cross section. As such, it enables a more comprehensive evaluation of white matter changes and has been successfully applied in other disease states^29^ ^41^ for more directly interpretable measures of WM connectivity.

With respects to limitations of the current study, direct comparisons between these two cohorts are limited by differences in MRI acquisition, participant number, age and study design between the two cohorts that required different statistical methodology. To investigate the influence of different acquisition parameters, we replicated our analysis in a subset of the TrackON-HD cohort who had additional multi-shell dMRI at the final time point with b-values of up to 2,000 s/mm². These results replicated the findings from the single-shell analyses and in addition revealed more widespread significant changes in FDC. This suggests improved sensitivity to WM differences in line with recent findings that higher b-values provide a more sensitive measure of FD^49^. This further strengthens findings in the HD-YAS cohort with a similar acquisition, that cortico-basal ganglia WM tracts are preserved. The HD-YAS cohort had fewer participants (n = 107), compared to the Track-ON HD study (n=157), however the young adult cohort is more difficult to recruit as the uptake of predictive genetic testing is much lower in this cohort^50^. In the Track-ON HD study there were significant age differences between preHD and controls, while this is an unavoidable consequence of the natural history of HD, we aimed to minimise this effect by including age as a covariate of no interest in all analyses.

The absence of significant difference between preHD and controls, 25 years before onset, does not exclude the possibility of subtle changes in cortico-basal ganglia WM tracts or the possibility of functional changes, which could be measured using functional MRI or magentoencephalography, and this warrants further investigation. Longitudinal follow up of this cohort will also be important pinpoint the exact time when WM loss begins.

## Conclusion

These findings suggest that WM tracts of cortico-basal ganglia functional sub-regions remain intact in preHD gene-carriers approximately 25 years before to predicted onset and that degeneration begins within an 11-25 year time frame from diagnosis. Selective vulnerability is seen in WM tracts to the limbic and motor striatum and the motor and sensory thalamus. This indicates that initiation of disease modifying therapies before demonstrable changes have occurred could prevent neurodegeneration of these WM tracts and highlights selectively vulnerable sub-regions of the striatum and thalamus that may be important targets for future therapies.

## Acknowledgements

The authors wish to extend their gratitude to the study participants and their families who supported them. Thanks also to the staff at the Wellcome Centre for Human Neuroimaging.

## Author contributions

PZ; 1) conception and design of the study, 2) acquisition and analysis of data or 3) drafting a significant portion of the manuscript and figures, PM; 1) conception and design of the study, 2) analysis of data and 3) drafting a significant portion of the manuscript, SG; 2) acquisition and analysis of data, EBJ; 2) acquisition and analysis of data, MP; 2) acquisition and analysis of data, AN; 2) acquisition and analysis of data, RIS; 2) acquisition and analysis of data, GR; 1) conception and design of the study and SJT; 1) conception and design of the study.

## Full financial disclosure

PZ, SG, EBJ, MP, AN, RIS, GR and SJT were supported by grant funding from the Wellcome Trust (ref. 200181/Z/15/Z) AN also received support from the Leonard Wolfson Foundation. PM is funded by the National Institute for Health Research. TH is funded by the Murdoch Children’s Research Institute (Melbourne) from the 1^st^ of January 2020, and prior to that from the National Health and Medical Research Council (NHMRC) of Australia. The authors report no conflicts of interest.

GR reports grants from Wellcome Trust, during the conduct of the study and outside of the current study, the MRC, UKRI and NIHR; and consultancy fees from Google Health. SJT receives grant funding for her research from the Medical Research Council UK, the Wellcome Trust, the Rosetrees Trust, Takeda Pharmaceuticals Ltd, Vertex Pharmaceuticals, Cantervale Limited, NIHR North Thames Local Clinical Research Network, UK Dementia Research Institute, and the CHDI Foundation. In the past 2 years, SJT has undertaken consultancy services, including advisory boards, with Alnylam Pharmaceuticals Inc., Annexon Inc., DDF Discovery Ltd, F. Hoffmann-La Roche Ltd, Genentech, PTC Bio, Novartis Pharma, Takeda Pharmaceuticals Ltd, Triplet Therapeutics, UCB Pharma S.A., University College Irvine and Vertex Pharmaceuticals Incorporated. All honoraria for these consultancies were paid through the offices of UCL Consultants Ltd., a wholly owned subsidiary of University College London.

## Working Group Authors

HD-YAS Investigators are K Osborne-Crowley, C Parker, J Lowe, C Estevez-Fraga, K Fayer, H Wellington, FB Rodrigues, LM Byrne, A Heselgrave, H Hyare, H Zetterberg, EJ Wild, H Zhang (University College London), C O’Callaghan, Christelle Langley, TW Robbins, BJ Sahakian (University of Cambridge), C Sampaio (CHDI Management/CHDI Foundation Inc), D Langbehn (University of Iowa).

TrackOn-HD Investigators are BR Leavitt, A Coleman, J Decolongon, M Fan, T Petkau, Y Koren (University of British Columbia, Vancouver); A Durr, C Jauffret, D Justo, S Lehericy, K Nigaud, R Valabrègue, (ICM and APHP, Pitié-Salpêtrière University Hospital, Paris). RAC Roos, A Schoonderbeek, E P ‘t Hart, S van den Bogaard, (Leiden University Medical Centre, Leiden); A Razi, R Ghosh, DJ Hensman Moss, H Crawford, C Berna,, G Owen, I Malone, J Read (University College London, London). R Reilmann, N Weber (George Huntington Institute, Munster); J Stout, S Andrews, A O’Regan, I Labuschagne, (Monash University, Melbourne); B Landwehrmeyer, M Orth, I Mayer (University of Ulm, Ulm); H Johnson, D Langbehn, J Long, J Mills (University of Iowa); A Cassidy, C Frost, R Keogh (London School of Hygiene and Tropical Medicine, London); B Borowsky (CHDI, USA); D Craufurd (University of Manchester); E Scheller, S Kloppel, L Mincova (Freiburg University).

## References

1. Ross CA, Aylward EH, Wild EJ, et al. Huntington disease: natural history, biomarkers and prospects for therapeutics. Nat Rev Neurol 2014;10(4):204–16. doi: 10.1038/nrneurol.2014.24

2. Shepherd GM. Corticostriatal connectivity and its role in disease. Nat Rev Neurosci 2013;14(4):278–91. doi: 10.1038/nrn3469

3. Tabrizi SJ, Scahill RI, Durr A, et al. Biological and clinical changes in premanifest and early stage Huntington’s disease in the TRACK-HD study: the 12-month longitudinal analysis. Lancet Neurol 2011;10(1):31–42. doi: 10.1016/S1474-4422(10)70276-3 [published Online First: 2010/12/07]

4. McColgan P, Seunarine KK, Razi A, et al. Selective vulnerability of Rich Club brain regions is an organizational principle of structural connectivity loss in Huntington’s disease. Brain 2015;138(Pt 11):3327–44. doi: 10.1093/brain/awv259

5. McColgan P, Seunarine KK, Gregory S, et al. Topological length of white matter connections predicts their rate of atrophy in premanifest Huntington’s disease. JCI Insight 2017;2(8) doi: 10.1172/jci.insight.92641

6. Zhang J, Gregory S, Scahill RI, et al. In vivo characterization of white matter pathology in premanifest huntington’s disease. Ann Neurol 2018;84(4):497–504. doi: 10.1002/ana.25309

7. Bohanna I, Georgiou-Karistianis N, Egan GF. Connectivity-based segmentation of the striatum in Huntington’s disease: vulnerability of motor pathways. Neurobiol Dis 2011;42(3):475–81. doi: 10.1016/j.nbd.2011.02.010 [published Online First: 2011/03/09]

8. Marrakchi-Kacem L, Delmaire C, Guevara P, et al. Mapping Cortico-Striatal Connectivity onto the Cortical Surface: A New Tractography-Based Approach to Study Huntington Disease. PLoS One 2013;8(2):e53135. doi: 10.1371/journal.pone.0053135 [published Online First: 2013/02/14]

9. Douaud G, Behrens TE, Poupon C, et al. In vivo evidence for the selective subcortical degeneration in Huntington’s disease. Neuroimage 2009;46(4):958–66. doi: 10.1016/j.neuroimage.2009.03.044 [published Online First: 2009/04/01]

10. Kloppel S, Draganski B, Golding CV, et al. White matter connections reflect changes in voluntary-guided saccades in pre-symptomatic Huntington’s disease. Brain 2008;131(Pt 1):196–204. doi: 10.1093/brain/awm275 [published Online First: 2007/12/07]

11. Shaffer JJ, Ghayoor A, Long JD, et al. Longitudinal diffusion changes in prodromal and early HD: Evidence of white-matter tract deterioration. Hum Brain Mapp 2017;38(3):1460–77. doi: 10.1002/hbm.23465

12. Tabrizi SJ, Flower MD, Ross CA, et al. Huntington disease: new insights into molecular pathogenesis and therapeutic opportunities. Nat Rev Neurol 2020 doi: 10.1038/s41582-020-0389-4 [published Online First: 2020/08/17]

13. Kloppel S, Gregory S, Scheller E, et al. Compensation in Preclinical Huntington’s Disease: Evidence From the Track-On HD Study. EBioMedicine 2015;2(10):1420–9. doi: 10.1016/j.ebiom.2015.08.002

14. Scahill RI, Zeun P, Osborne-Crowley K, et al. Biological and clinical characteristics of gene carriers far from predicted onset in the Huntington’s disease Young Adult Study (HD-YAS): a cross-sectional analysis. Lancet Neurol 2020;19(6):502–12. doi: 10.1016/S1474-4422(20)30143-5

15. Langbehn DR, Brinkman RR, Falush D, et al. A new model for prediction of the age of onset and penetrance for Huntington’s disease based on CAG length. Clin Genet 2004;65(4):267–77. doi: 10.1111/j.1399-0004.2004.00241.x [published Online First: 2004/03/18]

16. Scahill RI, Zeun P, Osborne-Crowley K, et al. Biological and clinical characteristics of gene carriers far from predicted onset in the Huntington’s disease Young Adult Study (HD-YAS): a cross-sectional analysis. The Lancet Neurology 2020 doi: In press [published Online First: In press]

17. Penney JB, Jr., Vonsattel JP, MacDonald ME, et al. CAG repeat number governs the development rate of pathology in Huntington’s disease. Annals of neurology 1997;41(5):689–92. doi: 10.1002/ana.410410521 [published Online First: 1997/05/01]

18. Kloppel S, Gregory S, Scheller E, et al. Compensation in Preclinical Huntington’s Disease: Evidence From the Track-On HD Study. EBioMedicine 2015;2(10):1420–9. doi: 10.1016/j.ebiom.2015.08.002 [published Online First: 2015/12/03]

19. Tournier JD, Smith R, Raffelt D, et al. MRtrix3: A fast, flexible and open software framework for medical image processing and visualisation. Neuroimage 2019;202:116137. doi: 10.1016/j.neuroimage.2019.116137

20. Jenkinson M, Beckmann CF, Behrens TE, et al. Fsl. Neuroimage 2012;62(2):782–90. doi: 10.1016/j.neuroimage.2011.09.015

21. Veraart J, Novikov DS, Christiaens D, et al. Denoising of diffusion MRI using random matrix theory. Neuroimage 2016;142:394–406. doi: 10.1016/j.neuroimage.2016.08.016

22. Kellner E, Dhital B, Kiselev VG, et al. Gibbs-ringing artifact removal based on local subvoxel-shifts. Magn Reson Med 2016;76(5):1574–81. doi: 10.1002/mrm.26054

23. Andersson JL, Sotiropoulos SN. An integrated approach to correction for off-resonance effects and subject movement in diffusion MR imaging. Neuroimage 2016;125:1063–78. doi: 10.1016/j.neuroimage.2015.10.019

24. Dyrby TB, Lundell H, Burke MW, et al. Interpolation of diffusion weighted imaging datasets. Neuroimage 2014;103:202–13. doi: 10.1016/j.neuroimage.2014.09.005

25. Raffelt DA, Tournier JD, Smith RE, et al. Investigating white matter fibre density and morphology using fixel-based analysis. Neuroimage 2017;144(Pt A):58–73. doi: 10.1016/j.neuroimage.2016.09.029

26. Dhollander T, Raffelt D, Connelly A. Unsupervised 3-tissue response function estimation from single-shell or multi-shell diffusion MR data without a co-registered T1 image2016.

27. Dhollander T, Connelly A. A novel iterative approach to reap the benefits of multi-tissue CSD from just single-shell (+b=0) diffusion MRI data2016.

28. Jeurissen B, Tournier JD, Dhollander T, et al. Multi-tissue constrained spherical deconvolution for improved analysis of multi-shell diffusion MRI data. Neuroimage 2014;103:411–26. doi: 10.1016/j.neuroimage.2014.07.061

29. Mito R, Raffelt D, Dhollander T, et al. Fibre-specific white matter reductions in Alzheimer’s disease and mild cognitive impairment. Brain 2018;141(3):888–902. doi: 10.1093/brain/awx355

30. Raffelt D, Tournier JD, Fripp J, et al. Symmetric diffeomorphic registration of fibre orientation distributions. Neuroimage 2011;56(3):1171–80. doi: 10.1016/j.neuroimage.2011.02.014

31. Raffelt D, Tournier JD, Rose S, et al. Apparent Fibre Density: a novel measure for the analysis of diffusion-weighted magnetic resonance images. Neuroimage 2012;59(4):3976–94. doi: 10.1016/j.neuroimage.2011.10.045

32. Tziortzi AC, Haber SN, Searle GE, et al. Connectivity-based functional analysis of dopamine release in the striatum using diffusion-weighted MRI and positron emission tomography. Cereb Cortex 2014;24(5):1165–77. doi: 10.1093/cercor/bhs397

33. Behrens TE, Johansen-Berg H, Woolrich MW, et al. Non-invasive mapping of connections between human thalamus and cortex using diffusion imaging. Nat Neurosci 2003;6(7):750–7. doi: 10.1038/nn1075

34. Modat M, Ridgway GR, Taylor ZA, et al. Fast free-form deformation using graphics processing units. Comput Methods Programs Biomed 2010;98(3):278–84. doi: 10.1016/j.cmpb.2009.09.002 [published Online First: 2009/10/13]

35. Zalesky A, Fornito A. A DTI-derived measure of cortico-cortical connectivity. IEEE Trans Med Imaging 2009;28(7):1023–36. doi: 10.1109/TMI.2008.2012113

36. Harrington DL, Long JD, Durgerian S, et al. Cross-sectional and longitudinal multimodal structural imaging in prodromal Huntington’s disease. Mov Disord 2016 doi: 10.1002/mds.26803

37. Chatterjee A, Anderson KE, Moskowitz CB, et al. A comparison of self-report and caregiver assessment of depression, apathy, and irritability in Huntington’s disease. J Neuropsychiatry Clin Neurosci 2005;17(3):378–83. doi: 10.1176/appi.neuropsych.17.3.378

38. Huntington Study Group CI. Unified Huntington’s Disease Rating Scale: reliability and consistency. Mov Disord 1996;11(2):136–42. doi: 10.1002/mds.870110204

39. Barnat M, Le Friec J, Benstaali C, et al. Huntingtin-Mediated Multipolar-Bipolar Transition of Newborn Cortical Neurons Is Critical for Their Postnatal Neuronal Morphology. Neuron 2017;93(1):99–114. doi: 10.1016/j.neuron.2016.11.035

40. Barnat M, Capizzi M, Aparicio E, et al. Huntington’s disease alters human neurodevelopment. Science 2020;369(6505):787–93. doi: 10.1126/science.aax3338

41. Zarkali A, McColgan P, Leyland LA, et al. Fiber-specific white matter reductions in Parkinson hallucinations and visual dysfunction. Neurology 2020;94(14):e1525–e38. doi: 10.1212/WNL.0000000000009014

42. Tabrizi SJ, Ghosh R, Leavitt BR. Huntingtin Lowering Strategies for Disease Modification in Huntington’s Disease. Neuron 2019;101(5):801–19. doi: 10.1016/j.neuron.2019.01.039

43. Evers MM, Miniarikova J, Juhas S, et al. AAV5-miHTT Gene Therapy Demonstrates Broad Distribution and Strong Human Mutant Huntingtin Lowering in a Huntington’s Disease Minipig Model. Mol Ther 2018;26(9):2163–77. doi: 10.1016/j.ymthe.2018.06.021 [published Online First: 2018/07/17]

44. Rub U, Seidel K, Heinsen H, et al. Huntington’s disease (HD): the neuropathology of a multisystem neurodegenerative disorder of the human brain. Brain Pathol 2016;26(6):726–40. doi: 10.1111/bpa.12426

45. Poudel GR, Stout JC, Dominguez DJ, et al. Longitudinal change in white matter microstructure in Huntington’s disease: The IMAGE-HD study. Neurobiology of disease 2015;74:406–12. doi: 10.1016/j.nbd.2014.12.009 [published Online First: 2014/12/17]

46. McColgan P, Razi A, Gregory S, et al. Structural and functional brain network correlates of depressive symptoms in premanifest Huntington’s disease. Hum Brain Mapp 2017 doi: 10.1002/hbm.23527

47. Jones DK, Knosche TR, Turner R. White matter integrity, fiber count, and other fallacies: the do’s and don’ts of diffusion MRI. Neuroimage 2013;73:239–54. doi: 10.1016/j.neuroimage.2012.06.081

48. Novak MJ, Seunarine KK, Gibbard CR, et al. White matter integrity in premanifest and early Huntington’s disease is related to caudate loss and disease progression. Cortex 2014;52:98–112. doi: 10.1016/j.cortex.2013.11.009

49. Genc S, Tax CMW, Raven EP, et al. Impact of b-value on estimates of apparent fibre density. Human brain mapping 2020 doi: 10.1002/hbm.24964 [published Online First: 2020/03/28]

50. Baig SS, Strong M, Rosser E, et al. 22 Years of predictive testing for Huntington’s disease: the experience of the UK Huntington’s Prediction Consortium. Eur J Hum Genet 2016;24(10):1396–402. doi: 10.1038/ejhg.2016.36 [published Online First: 2016/05/12]

