## Supplemental tables for "Timing of selective basal ganglia white matter loss in Huntington’s disease"

**Supplementary Tables**

**S1. Missing data**

| Study | QC Fail | Left handed or ambidexterous | Total |
| --- | --- | --- | --- |
| HD-YAS  (N 122) | 1 | 14 | 107 |
| TrackOn-HD Single-shell 2012 (N=222) | 41 | 24 | 157 |
| TrackOn-HD Single-shell 2013 (N=222) | 30 | 24 | 168 |
| TrackOn-HD Single-shell 2014 (N=222) | 40 | 24 | 158 |
| TrackOn-HD Multi-shell (N=80) | 0 | 7 | 73 |

For TrackOn-HD single shell data, 56 gene carriers and 65 controls had data at 3 time points, 28 premanifest and 24 controls had data at 2-time points, and 10 gene carriers and 9 controls had data at one time point. The TrackOn-HD multi-shell acquisition was only performed at 2 of the 4 sites (London and Paris).

### S2. MRI acquisitions and demographics

|  | HD-YAS | | | TrackON-HD  Single-Shell | | |  | TrackON-HD Multi-Shell |  |
| --- | --- | --- | --- | --- | --- | --- | --- | --- | --- |
|  | **PreHD**  **N=54** | **Control**  **N=53** | **p** | **PreHD**  **N=72** | **Control**  **N=85** | **p** | **PreHD**  **N=33** | **Control**  **N=40** | **p** |
| Age | 29.8 ± 5.6 | 29.3 ± 5.5 | 0.67 | 43.3 ± 9.2 | 48.8 ± 9.8 | 4.0 x 10^-4^ | 42.1 ± 9.5 | 47.2 ± 10.6 | 0.03 |
| Male (%) | 48 | 43 | 0.49 | 53 | 38 | 0.06 | 42 | 63 | 0.09 |
| Education | 4.9 ± 1.8 | 4.8 ± 1.6 | 0.71 | 4.0 ± 1.0 | 4.0 ± 1.0 | 0.94 | 3.9 ± 1.2 | 4.0 ± 0.9 | 0.28 |
| CAG | 42.2 ± 1.7 |  |  | 42.9 ± 2.3 |  |  | 43.0 ± 1.9 |  |  |
| Years to Onset | 24.7 ± 6.3 |  |  | 11.3 ± 3.9 |  |  | 11.5 ± 4.0 |  |  |
| MRI | Prisma, Siemens | | | Tim Trio, Siemens  Achieva, Phillips | | | Tim Trio, Siemens | | |
| b-Value | 0, 300, 1000, 2000 | | | 0, 1000 | | | 0, 300, 700, 2000 | | |
| Gradient Directions | 10, 8, 64, 64 | | | 8, 42 (Siemens)  1, 42 (Phillips) | | | 14, 8, 32, 64 | | |
| Voxel-size (mm) | 2 x 2 x 2 | | | 2 x 2 x 2 (Siemens)  1.96 x 1.96 x 1.96 (Philips) | | | 2.5 x 2.5 x 2.5 | | |
| TR/TE (ms) | 3260/58 | | | 13100/88 (Siemens)  11000/56 (Philips) | | | 7000 /90.8 | | |
| Slices | 72 | | | 75 | | | 55 | | |
| Acquisition time | 15 mins | | | 10 mins | | | 15 mins | | |

Unless otherwise specified, values are means ± standard deviations. Group comparisons were made using t tests (age, education) or chi-squared (sex and education). Education was measured using international standard classification of education (ISCED). TrackON-HD demographics presented for baseline visit. All MRI are 3 tesla. TR=repetition time, TE=echo time.

S3. Cortico-striatal FDC in TrackON-HD Single-Shell Longitudinal

| Cortico-striatal Tract | *δ* | SE | p | FDR |
| --- | --- | --- | --- | --- |
| L Limbic | -0.002 | 0.003 | 0.52 | 0.60 |
| R Limbic | -0.003 | 0.002 | 0.26 | 0.45 |
| L Cognitive | -0.002 | 0.001 | 0.20 | 0.36 |
| R Cognitive | -0.002 | 0.001 | 0.25 | 0.45 |
| L Rostral Motor | -0.003 | 0.002 | 0.14 | 0.33 |
| R Rostral Motor | -0.001 | 0.002 | 0.70 | 0.82 |
| L Caudal Motor | -0.004 | 0.002 | 0.09 | 0.30 |
| R Caudal Motor | 0.002 | 0.002 | 0.42 | 0.58 |
| L Parietal | -0.002 | 0.002 | 0.37 | 0.51 |
| R Parietal | 0.001 | 0.002 | 0.82 | 0.82 |
| L Temporal | 3.0 x 10^-4^ | 0.002 | 0.80 | 0.89 |
| R Temporal | -0.003 | 0.002 | 0.12 | 0.45 |
| L Occipital | -0.003 | 0.002 | 0.08 | 0.30 |
| R Occipital | -0.003 | 0.002 | 0.17 | 0.45 |

### Unadjusted means displayed. *δ*=estimated group slope difference, SE=standard error, p = p-value, FDR = false discovery rate corrected p-value.

S4. Cortico-thalamic FDC in TrackON-HD Single-Shell Longitudinal

| Cortico-thalamic Tract | *δ* | SE | p | FDR |
| --- | --- | --- | --- | --- |
| L Prefrontal | 0.000 | 0.002 | 0.81 | 0.81 |
| R Prefrontal | -0.001 | 0.001 | 0.64 | 0.75 |
| L Premotor | -0.002 | 0.002 | 0.40 | 0.48 |
| R Premotor | 0.002 | 0.002 | 0.39 | 0.75 |
| L Primary Motor | -0.002 | 0.003 | 0.37 | 0.48 |
| R Primary Motor | 0.001 | 0.002 | 0.59 | 0.75 |
| L Sensory | -0.002 | 0.002 | 0.41 | 0.48 |
| R Sensory | 0.000 | 0.002 | 0.95 | 0.95 |
| L Parietal | -0.001 | 0.002 | 0.34 | 0.48 |
| R Parietal | -0.002 | 0.002 | 0.28 | 0.75 |
| L Temporal | -0.001 | 0.001 | 0.32 | 0.48 |
| R Temporal | -0.001 | 0.001 | 0.44 | 0.75 |
| L Occipital | -0.004 | 0.002 | 0.03 | 0.24 |
| R Occipital | -0.004 | 0.002 | 0.04 | 0.28 |

### Unadjusted means displayed. *δ*=estimated group slope difference, SE=standard error, p = p-value, FDR = false discovery rate corrected p-value.

S5. Cortico-striatal FDC TrackON-HD Multi-Shell

| **Cortico-striatal Tract** | **Control Mean** | **PreHD Mean** | **SE** | **p** | **FDR** |
| --- | --- | --- | --- | --- | --- |
| **L Limbic** | 0.39 | 0.38 | 0.001 | 0.008 | 0.01 |
| **R Limbic** | 0.41 | 0.40 | 0.001 | 0.005 | 0.01 |
| **L Cognitive** | 0.43 | 0.42 | 0.001 | 0.02 | 0.02 |
| **R Cognitive** | 0.43 | 0.42 | 0.001 | 0.002 | 0.01 |
| **L Rostral Motor** | 0.50 | 0.50 | 0.001 | 0.04 | 0.05 |
| **R Rostral Motor** | 0.51 | 0.51 | 0.001 | 0.19 | 0.19 |
| **L Caudal Motor** | 0.59 | 0.55 | 0.001 | 0.002 | 0.01 |
| **R Caudal Motor** | 0.60 | 0.58 | 0.001 | 0.02 | 0.02 |
| **L Parietal** | 0.58 | 0.56 | 0.001 | 0.005 | 0.01 |
| **R Parietal** | 0.56 | 0.55 | 0.001 | 0.01 | 0.02 |
| **L Temporal** | 0.41 | 0.40 | 0.001 | 0.01 | 0.10 |
| **R Temporal** | 0.41 | 0.40 | 0.001 | 0.04 | 0.05 |
| **L Occipital** | 0.58 | 0.57 | 0.001 | 0.008 | 0.01 |
| **R Occipital** | 0.59 | 0.56 | 0.001 | 0.006 | 0.01 |

### Unadjusted means displayed. γ=estimated group intercept difference, SE=standard error, p = p-value, FDR = false discovery rate corrected p-value.

S6. Cortico-thalamic FDC TrackON-HD Multi-Shell

| Cortico-thalamic Tract | Control Mean | PreHD Mean | SE | p | FDR |
| --- | --- | --- | --- | --- | --- |
| L Prefrontal | 0.49 | 0.47 | 0.001 | 0.001 | 0.01 |
| R Prefrontal | 0.54 | 0.52 | 0.001 | 0.006 | 0.01 |
| L Premotor | 0.58 | 0.54 | 0.001 | 3.0 x 10^-4^ | 0.01 |
| R Premotor | 0.56 | 0.54 | 0.001 | 0.04 | 0.05 |
| L Primary Motor | 0.49 | 0.47 | 0.001 | 0.01 | 0.02 |
| R Primary Motor | 0.45 | 0.43 | 0.001 | 0.02 | 0.03 |
| L Sensory | 0.46 | 0.45 | 0.001 | 0.06 | 0.07 |
| R Sensory | 0.44 | 0.42 | 0.001 | 0.007 | 0.01 |
| L Parietal | 0.50 | 0.48 | 0.001 | 0.006 | 0.01 |
| R Parietal | 0.52 | 0.49 | 0.001 | 0.001 | 0.01 |
| L Temporal | 0.52 | 0.51 | 0.001 | 0.02 | 0.02 |
| R Temporal | 0.53 | 0.50 | 0.001 | 0.002 | 0.01 |
| L Occipital | 0.50 | 0.48 | 0.001 | 0.04 | 0.05 |
| R Occipital | 0.44 | 0.41 | 0.001 | 0.001 | 0.01 |

### Unadjusted means displayed. γ=estimated group intercept difference, SE=standard error, p = p-value, FDR = false discovery rate corrected p-value.

### S7. Cortico-striatal fibre density (FD) and fibre cross-section (FC) in HD-YAS

| Cortico-Striatal Tract | Control Mean | PreHD Mean | SE | p | FDR |
| --- | --- | --- | --- | --- | --- |
| L Limbic FD | 0.50 | 0.50 | 2.3 x 10^-4^ | 0.44 | 0.57 |
| L Limbic FC | 0.07 | 0.06 | 8.5 x 10^-4^ | 0.20 | 0.49 |
| R Limbic FD | 0.49 | 0.49 | 2.1 x 10^-4^ | 0.19 | 0.49 |
| R Limbic FC | 0.08 | 0.07 | 8.8 x 10^-4^ | 0.17 | 0.49 |
| L Cognitive FD | 0.54 | 0.54 | 2.0 x 10^-4^ | 0.37 | 0.52 |
| L Cognitive FC | 0.06 | 0.06 | 9.3 x 10^-4^ | 0.33 | 0.50 |
| R Cognitive FD | 0.53 | 0.53 | 1.9 x 10^-4^ | 0.51 | 0.59 |
| R Cognitive FC | 0.05 | 0.06 | 8.6 x 10^-4^ | 0.47 | 0.59 |
| L Rostral Motor FD | 0.55 | 0.55 | 2.2 x 10^-4^ | 0.14 | 0.49 |
| L Rostral Motor FC | 0.04 | 0.06 | 9.2 x 10^-4^ | 0.78 | 0.79 |
| R Rostral Motor FD | 0.56 | 0.55 | 2.3 x 10^-4^ | 0.36 | 0.51 |
| R Rostral Motor FC | 0.05 | 0.05 | 8.2 x 10^-4^ | 0.32 | 0.50 |
| L Caudal Motor FD | 0.58 | 0.58 | 2.3 x 10^-4^ | 0.48 | 0.59 |
| L Caudal Motor FC | 0.06 | 0.03 | 8.9 x 10^-4^ | 0.03 | 0.48 |
| R Caudal Motor FD | 0.58 | 0.58 | 2.4 x 10^-4^ | 0.73 | 0.75 |
| R Caudal Motor FC | 0.05 | 0.03 | 7.7 x 10^-4^ | 0.08 | 0.48 |
| L Parietal FD | 0.66 | 0.66 | 3.0 x 10^-4^ | 0.42 | 0.55 |
| L Parietal FC | 0.01 | 0.01 | 8.1 x 10^-4^ | 0.38 | 0.52 |
| R Parietal FD | 0.67 | 0.67 | 3.1 x 10^-4^ | 0.55 | 0.60 |
| R Parietal FC | 0.01 | 0.01 | 8.1 x 10^-4^ | 0.28 | 0.50 |
| L Temporal FD | 0.56 | 0.56 | 2.4 x 10^-4^ | 0.63 | 0.67 |
| L Temporal FC | 0.05 | 0.04 | 7.5 x 10^-4^ | 0.10 | 0.48 |
| R Temporal FD | 0.59 | 0.58 | 2.7 x 10^-4^ | 0.03 | 0.48 |
| R Temporal FC | 0.07 | 0.05 | 7.0 x 10^-4^ | 0.06 | 0.48 |
| L Occipital FD | 0.65 | 0.65 | 2.2 x 10^-4^ | 0.57 | 0.63 |
| L Occipital FC | 0.11 | 0.09 | 7.1 x 10^-4^ | 0.04 | 0.48 |
| R Occipital FD | 0.65 | 0.65 | 2.3 x 10^-4^ | 0.32 | 0.50 |
| R Occipital FC | 0.08 | 0.07 | 6.7 x 10^-4^ | 0.21 | 0.49 |

### S8. Cortico-thalamic fibre density (FD) and fibre cross-section (FC) in HD-YAS

| Cortico-thalamic Tract | Control Mean | PreHD Mean | SE | p | FDR |
| --- | --- | --- | --- | --- | --- |
| L Prefrontal FD | 0.59 | 0.59 | 2.2 x 10^-4^ | 0.17 | 0.49 |
| L Prefrontal FC | 0.02 | 0.01 | 7.9 x 10^-4^ | 0.25 | 0.50 |
| R Prefrontal FD | 0.6 | 0.6 | 2.3 x 10^-4^ | 0.28 | 0.50 |
| R Prefrontal FC | 0.02 | 0.02 | 7.6 x 10^-4^ | 0.38 | 0.52 |
| L Premotor FD | 0.62 | 0.62 | 2.5 x 10^-4^ | 0.53 | 0.60 |
| L Premotor FC | 0.04 | 0.02 | 8.2 x 10^-4^ | 0.05 | 0.48 |
| R Premotor FD | 0.6 | 0.61 | 2.5 x 10^-4^ | 0.81 | 0.81 |
| R Premotor FC | 0.03 | 0.03 | 7.8 x 10^-4^ | 0.19 | 0.49 |
| L Primary Motor FD | 0.59 | 0.6 | 2.6 x 10^-4^ | 0.62 | 0.66 |
| L Primary Motor FC | 0.02 | 0.02 | 7.9 x 10^-4^ | 0.27 | 0.50 |
| R Primary Motor FD | 0.58 | 0.58 | 2.7 x 10^-4^ | 0.69 | 0.72 |
| R Primary Motor FC | 0.02 | 0.02 | 8.5 x 10^-4^ | 0.26 | 0.50 |
| L Sensory FD | 0.61 | 0.61 | 3.0 x 10^-4^ | 0.51 | 0.59 |
| L Sensory FC | 0.02 | 0.02 | 8.3 x 10^-4^ | 0.49 | 0.59 |
| R Sensory FD | 0.56 | 0.56 | 3.2 x 10^-4^ | 0.46 | 0.59 |
| R Sensory FC | 0.03 | 0.03 | 9.0 x 10^-4^ | 0.26 | 0.50 |
| L Parietal FD | 0.64 | 0.64 | 2.5 x 10^-4^ | 0.21 | 0.49 |
| L Parietal FC | 0.06 | 0.05 | 7.2 x 10^-4^ | 0.15 | 0.49 |
| R Parietal FD | 0.62 | 0.61 | 2.6 x 10^-4^ | 0.12 | 0.49 |
| R Parietal FC | 0.03 | 0.04 | 6.6 x 10^-4^ | 0.36 | 0.51 |
| L Temporal FD | 0.51 | 0.5 | 3.0 x 10^-4^ | 0.42 | 0.55 |
| L Temporal FC | 0.01 | 3.0 x 10^-4^ | 8.3 x 10^-4^ | 0.10 | 0.48 |
| R Temporal FD | 0.51 | 0.49 | 3.4 x 10^-4^ | 0.05 | 0.48 |
| R Temporal FC | 0.001 | 0.007 | 8.0 x 10^-4^ | 0.55 | 0.60 |
| L Occipital FD | 0.47 | 0.46 | 1.4 x 10^-4^ | 0.04 | 0.48 |
| L Occipital FC | 0.08 | 0.08 | 7.2 x 10^-4^ | 0.23 | 0.50 |
| R Occipital FD | 0.51 | 0.5 | 1.6 x 10^-4^ | 0.10 | 0.48 |
| R Occipital FC | 0.07 | 0.07 | 7.2 x 10^-4^ | 0.32 | 0.50 |

### S9. Cortico-striatal fibre density (FD) and fibre cross-section (FC) in TrackON-HD Single-Shell Baseline

| Cortico-Striatal Tract | Control Mean | PreHD Mean | *γ* | SE | p | FDR |
| --- | --- | --- | --- | --- | --- | --- |
| L Limbic FD | 0.38 | 0.38 | -0.005 | 0.004 | 0.28 | 0.39 |
| L Limbic FC | 0.09 | 0.06 | -0.06 | 0.01 | 1.2 x 10^-7^ | 1.6 x 10^-5^ |
| R Limbic FD | 0.36 | 0.36 | 1.7 x 10^-4^ | 0.005 | 0.97 | 0.97 |
| R Limbic FC | 0.10 | 0.07 | -0.06 | 0.01 | 4.9 x 10^-7^ | 6.9 x10^-6^ |
| L Cognitive FD | 0.40 | 0.41 | -0.001 | 0.003 | 0.79 | 0.79 |
| L Cognitive FC | 0.03 | 0.03 | -0.03 | 0.01 | 0.01 | 0.02 |
| R Cognitive FD | 0.40 | 0.41 | 0.003 | 0.003 | 0.35 | 0.46 |
| R Cognitive FC | 0.03 | 0.02 | -0.03 | 0.01 | 0.006 | 0.01 |
| L Rostral Motor FD | 0.46 | 0.46 | 0.003 | 0.004 | 0.37 | 0.44 |
| L Rostral Motor FC | 0.07 | 0.07 | -0.03 | 0.01 | 0.02 | 0.04 |
| R Rostral Motor FD | 0.47 | 0.47 | 0.002 | 0.003 | 0.59 | 0.63 |
| R Rostral Motor FC | 0.10 | 0.08 | -0.04 | 0.01 | 0.001 | 0.003 |
| L Caudal Motor FD | 0.50 | 0.50 | -0.003 | 0.004 | 0.37 | 0.44 |
| L Caudal Motor FC | 0.09 | 0.06 | -0.04 | 0.01 | 8.9 x 10^-5^ | 6.3 x 10^-4^ |
| R Caudal Motor FD | 0.51 | 0.50 | -0.003 | 0.004 | 0.39 | 0.46 |
| R Caudal Motor FC | 0.09 | 0.06 | -0.04 | 0.01 | 0.001 | 0.003 |
| L Parietal FD | 0.53 | 0.53 | -0.002 | 0.004 | 0.67 | 0.73 |
| L Parietal FC | 0.04 | 0.03 | -0.03 | 0.01 | 0.008 | 0.02 |
| R Parietal FD | 0.54 | 0.53 | -0.004 | 0.004 | 0.40 | 0.46 |
| R Parietal FC | 0.05 | 0.05 | -0.02 | 0.01 | 0.10 | 0.18 |
| L Temporal FD | 0.40 | 0.41 | 0.01 | 0.004 | 3.9 x 10^-4^ | 0.001 |
| L Temporal FC | 0.01 | -0.01 | -0.04 | 0.01 | 1.3x10^-4^ | 6.5 x 10^-4^ |
| R Temporal FD | 0.42 | 0.44 | 0.01 | 0.004 | 0.001 | 0.003 |
| R Temporal FC | 0.02 | 0.01 | -0.03 | 0.01 | 0.004 | 0.009 |
| L Occipital FD | 0.52 | 0.52 | 0.008 | 0.003 | 0.007 | 0.02 |
| L Occipital FC | 0.05 | 0.05 | -0.03 | 0.01 | 0.009 | 0.02 |
| R Occipital FD | 0.52 | 0.53 | 0.003 | 0.003 | 0.35 | 0.46 |
| R Occipital FC | 0.05 | 0.05 | -0.03 | 0.01 | 0.003 | 0.009 |

### S10. Cortico-thalamic fibre density (FD) and fibre cross-section (FC) in TrackON-HD Single-Shell Baseline

| Cortico-Thalamic Tract | Control Mean | PreHD Mean | *γ* | SE | p | FDR |
| --- | --- | --- | --- | --- | --- | --- |
| L Prefrontal FD | 0.47 | 0.47 | -0.003 | 0.003 | 0.32 | 0.35 |
| L Prefrontal FC | 0.02 | 0.01 | -0.03 | 0.01 | 0.02 | 0.06 |
| R Prefrontal FD | 0.47 | 0.47 | 0.001 | 0.003 | 0.75 | 0.79 |
| R Prefrontal FC | 0.02 | 0.01 | -0.03 | 0.01 | 0.02 | 0.06 |
| L Premotor FD | 0.52 | 0.51 | -0.008 | 0.004 | 0.06 | 0.08 |
| L Premotor FC | 0.09 | 0.07 | -0.03 | 0.01 | 0.006 | 0.02 |
| R Premotor FD | 0.51 | 0.51 | -0.007 | 0.004 | 0.08 | 0.16 |
| R Premotor FC | 0.09 | 0.08 | -0.03 | 0.01 | 0.02 | 0.06 |
| L Primary Motor FD | 0.53 | 0.52 | -0.009 | 0.004 | 0.05 | 0.08 |
| L Primary Motor FC | 0.08 | 0.05 | -0.04 | 0.01 | 3.3 x10^-4^ | 0.005 |
| R Primary Motor FD | 0.52 | 0.52 | -0.006 | 0.004 | 0.21 | 0.29 |
| R Primary Motor FC | 0.07 | 0.06 | -0.02 | 0.01 | 0.04 | 0.11 |
| L Sensory FD | 0.52 | 0.51 | -0.005 | 0.004 | 0.12 | 0.21 |
| L Sensory FC | 0.05 | 0.04 | -0.03 | 0.01 | 0.003 | 0.02 |
| R Sensory FD | 0.50 | 0.49 | -0.003 | 0.004 | 0.45 | 0.53 |
| R Sensory FC | 0.07 | 0.07 | -0.01 | 0.01 | 0.21 | 0.29 |
| L Parietal FD | 0.48 | 0.48 | 0.002 | 0.004 | 0.06 | 0.08 |
| L Parietal FC | 0.05 | 0.06 | -0.02 | 0.01 | 0.006 | 0.02 |
| R Parietal FD | 0.47 | 0.47 | -0.003 | 0.004 | 0.08 | 0.16 |
| R Parietal FC | 0.07 | 0.08 | -0.02 | 0.01 | 0.02 | 0.06 |
| L Temporal FD | 0.38 | 0.37 | -0.006 | 0.004 | 0.15 | 0.19 |
| L Temporal FC | -0.05 | -0.03 | -0.002 | 0.01 | 0.85 | 0.85 |
| R Temporal FD | 0.38 | 0.37 | -0.006 | 0.004 | 0.12 | 0.20 |
| R Temporal FC | -0.04 | -0.03 | -0.008 | 0.01 | 0.45 | 0.53 |
| L Occipital FD | 0.47 | 0.48 | 0.007 | 0.003 | 0.03 | 0.08 |
| L Occipital FC | 0.09 | 0.10 | -0.02 | 0.01 | 0.05 | 0.08 |
| R Occipital FD | 0.48 | 0.48 | 0.001 | 0.004 | 0.79 | 0.79 |
| R Occipital FC | 0.12 | 0.11 | -0.03 | 0.01 | 0.01 | 0.06 |

### S11. Cortico-striatal fibre density (FD) and fibre cross-section (FC) in TrackON-HD Multi-shell

| Cortico-Striatal Tract | Control Mean | PreHD Mean | SE | p | FDR |
| --- | --- | --- | --- | --- | --- |
| L Limbic FD | 0.36 | 0.36 | 4.1 x 10^-4^ | 0.05 | 0.08 |
| L Limbic FC | 0.06 | 0.05 | 0.001 | 0.01 | 0.03 |
| R Limbic FD | 0.38 | 0.37 | 3.4 x 10^-4^ | 0.10 | 0.13 |
| R Limbic FC | 0.06 | 0.05 | 0.001 | 0.00 | 0.02 |
| L Cognitive FD | 0.41 | 0.41 | 3.12 x 10^-4^ | 0.11 | 0.14 |
| L Cognitive FC | 0.02 | 0.01 | 0.001 | 0.01 | 0.02 |
| R Cognitive FD | 0.42 | 0.42 | 3.9 x 10^-4^ | 0.30 | 0.31 |
| R Cognitive FC | 0.03 | -0.007 | 0.001 | 0.00 | 0.01 |
| L Rostral Motor FD | 0.49 | 0.48 | 3.2 x 10^-4^ | 0.07 | 0.10 |
| L Rostral Motor FC | 0.02 | 0.01 | 0.001 | 0.08 | 0.11 |
| R Rostral Motor FD | 0.49 | 0.49 | 3.2 x 10^-4^ | 0.28 | 0.29 |
| R Rostral Motor FC | 0.03 | 0.03 | 0.001 | 0.19 | 0.22 |
| L Caudal Motor FD | 0.56 | 0.54 | 4.6 x 10^-4^ | 0.10 | 0.12 |
| L Caudal Motor FC | 0.04 | 1.6 | 0.001 | 0.01 | 0.03 |
| R Caudal Motor FD | 0.58 | 0.57 | 4.6 x 10^-4^ | 0.03 | 0.05 |
| R Caudal Motor FC | 0.02 | 0.01 | 0.001 | 0.16 | 0.20 |
| L Parietal FD | 0.58 | 0.57 | 4.2 x 10^-4^ | 0.07 | 0.10 |
| L Parietal FC | -0.009 | -0.02 | 0.001 | 0.02 | 0.04 |
| R Parietal FD | 0.556 | 0.55 | 4.4 x 10^-4^ | 0.06 | 0.09 |
| R Parietal FC | -0.001 | -0.001 | 0.001 | 0.08 | 0.10 |
| L Temporal FD | 0.36 | 0.36 | 3.9 x 10^-4^ | 0.27 | 0.28 |
| L Temporal FC | 0.1 | 0.09 | 0.001 | 0.03 | 0.06 |
| R Temporal FD | 0.37 | 0.37 | 3.6 x 10^-4^ | 0.44 | 0.44 |
| R Temporal FC | 0.09 | 0.07 | 0.001 | 0.01 | 0.02 |
| L Occipital FD | 0.53 | 0.53 | 3.6 x 10^-4^ | 0.46 | 0.46 |
| L Occipital FC | 0.09 | 0.05 | 0.001 | 0.00 | 0.01 |
| R Occipital FD | 0.52 | 0.52 | 4.3 x 10^-4^ | 0.23 | 0.25 |
| R Occipital FC | 0.1 | 0.06 | 0.001 | 0.00 | 0.01 |

### S12. Cortico-thalamic fibre density (FD) and fibre cross-section (FC) in TrackON-HD Multi-shell

| Cortico-thalamic Tract | Control Mean | PreHD Mean | SE | p | FDR |
| --- | --- | --- | --- | --- | --- |
| L Pre-frontal FD | 0.49 | 0.47 | 3 x 10^-4^ | 0.002 | 0.02 |
| L Pre-frontal FC | -0.002 | -0.02 | 0.001 | 0.01 | 0.03 |
| R Pre-frontal FD | 0.52 | 0.52 | 2.9 x 10^-4^ | 0.02 | 0.04 |
| R Pre-frontal FC | 0.01 | 1.9 | 0.001 | 0.02 | 0.04 |
| L Pre-motor FD | 0.56 | 0.54 | 4.4 x 10^-4^ | 0.01 | 0.02 |
| L Pre-motor FC | 0.01 | -0.02 | 0.001 | 0.01 | 0.02 |
| R Pre-motor FD | 0.55 | 0.54 | 4.6 x 10^-4^ | 0.05 | 0.09 |
| R Pre-motor FC | -0.001 | 0.007 | 0.001 | 0.27 | 0.28 |
| L Primary Motor FD | 0.5 | 0.48 | 4.1 x 10^-4^ | 0.06 | 0.10 |
| L Primary Motor FC | -0.02 | -0.03 | 0.010 | 0.05 | 0.07 |
| R Primary Motor FD | 0.45 | 0.44 | 3.8 x 10^-4^ | 0.02 | 0.04 |
| R Primary Motor FC | -0.02 | -0.03 | 0.001 | 0.07 | 0.10 |
| L Sensory Motor FD | 0.46 | 0.45 | 3.9 x 10^-4^ | 0.20 | 0.23 |
| L Sensory Motor FC | -0.01 | -0.01 | 0.002 | 0.12 | 0.14 |
| R Sensory Motor FD | 0.42 | 0.41 | 3.8 x 10^-4^ | 0.02 | 0.04 |
| R Sensory Motor FC | 0.01 | -0.02 | 0.001 | 0.02 | 0.04 |
| L Parietal FD | 0.47 | 0.47 | 3.5 x 10^-4^ | 0.25 | 0.27 |
| L Parietal FC | 0.05 | 0.02 | 0.001 | 0.004 | 0.02 |
| R Parietal FD | 0.48 | 0.48 | 3.7 x 10^-4^ | 0.22 | 0.24 |
| R Parietal FC | 0.06 | 0.02 | 0.001 | 5.0 x 10^-4^ | 0.01 |
| L Temporal FD | 0.49 | 0.49 | 6.0 x 10^-4^ | 0.27 | 0.28 |
| L Temporal FC | 0.04 | 0.02 | 0.001 | 0.02 | 0.04 |
| R Temporal FD | 0.5 | 0.49 | 4.9 x 10^-4^ | 0.09 | 0.12 |
| R Temporal FC | 0.04 | 0.003 | 0.001 | 0.003 | 0.02 |
| L Occipital FD | 0.44 | 0.44 | 4.1 x 10^-4^ | 0.21 | 0.23 |
| L Occipital FC | 0.11 | 0.09 | 0.001 | 0.004 | 0.02 |
| R Occipital FD | 0.37 | 0.36 | 3.1 x 10^-4^ | 0.09 | 0.12 |
| R Occipital FC | 0.16 | 0.12 | 0.001 | 2.00 | 0.01 |

**S13. Correlations between a priori cortico-striatal and cortico-thalamic FDC and corresponding clinical task**

| Correlation Track – Clinical measure | r | p |
| --- | --- | --- |
| L Caudal Motor – TMS | -0.22 | 0.007 |
| R Caudal Motor – TMS | -0.21 | 0.009 |
| L Limbic - apathy | 0.15 | 0.07 |
| R Limbic - apathy | 0.22 | 0.006 |
| L Pre-motor - TMS | -0.21 | 0.01 |
| R Pre-motor - TMS | -0.20 | 0.01 |
| L Primary Motor - TMS | -0.23 | 0.004 |
| R Primary Motor - TMS | -0.20 | 0.01 |

Correlations were performed using partial correlations with age, gender, and site as covariates. TMS = total motor score and DBS = disease burden score; derived from the Unified Huntington’s Disease Rating Scale (UHDRS). Apathy scores were from the Baltimore apathy/irritability scale. Caudal Motor and Limbic are cortico-striatal tracts. Premotor and Primary Motor are cortico-thalamic tracts.
